# Laminin N-terminus α31 is upregulated in invasive ductal breast cancer and changes the mode of tumour invasion

**DOI:** 10.1101/2020.05.28.120964

**Authors:** Lee D. Troughton, Tobias Zech, Kevin J. Hamill

**Author notes:** Corresponding author, Corresponding address: Institute of Life Course and Medical Sciences, William Henry Duncan Building, 6 West Derby Street, University of Liverpool, UK, L7 8TX.

## Abstract

Laminin N-terminus α31 (LaNt α31) is an alternative splice isoform derived from the laminin α3 gene. The LaNt α31 protein is enriched around the terminal duct lobular units in normal breast tissue. In the skin and cornea the protein influences epithelial cell migration and tissue remodelling. However, LaNt α31 has never been investigated in a tumour environment. Here we analysed LaNt α31 in invasive ductal carcinoma and determined its contribution to breast carcinoma invasion. LaNt α31 expression and distribution were analysed by immunohistochemistry in human breast tissue biopsy sections and tissue microarrays covering 232 breast cancer samples. This analysis revealed LaNt α31 to be upregulated in 56 % of invasive ductal carcinoma specimens compared with matched normal tissue, and further increased in nodal metastasis compared with the tumour mass in 45 % of samples. 65.8 % of triple negative cases displayed medium to high LaNt α31 expression. To study LaNt α31 function, an adenoviral system was used to induce expression in MCF-7 and MDA-MB-231 cells. Metabolic activity, 2D cell migration, and invasion into collagen hydrogels were not significantly different between LaNt α31 overexpressing cells and control treated cells. However, LaNt α31 overexpressing MDA-MB-231 cells displayed a striking change in their mode of invasion into laminin-containing Matrigel; changing from multicellular streaming to individual cellular-invasion. In agreement with these results, 66.7% of the tumours with the highest LaNt α31 expression were non-cohesive. Together these findings indicate that breast cancer-associated changes in LaNt α31 expression could directly contribute to tumour invasiveness, and that this little-studied protein may become a therapeutic target.

## Introduction

An essential stage of tumour progression is acquisition of an ability to breakthrough an organised extracellular matrix (ECM) structure termed the basement membrane (BM)[1]. Determining the expression and distribution of BM proteins has yielded valuable biomarkers to predict breast cancer outcomes [2–6]. Much of this work has focused on the laminin (LM) family of BM proteins, which are not only essential barrier components, but also act as substrates for tumour cell migration, regulate actin dynamics, influence survival and growth signalling pathways, and maintain quiescence in cancer stem cell niches; all of which functions influence breast cancer progression [7–10]. Here we investigated a relatively unstudied LM-related protein, laminin N-terminus α31 (LaNt α31) that we predicted would change in cancer and which could therefore represent a new target for therapeutic development [11, 12].

LMs are obligate heterotrimeric proteins comprised of an α, β and γ chain, with each chain derived from one of five α genes (LAMA1-5), one of three β (LAMB1-3), and one of three γ (LAMC1-3), as reviewed in [7, 8, 13]. Through the use of distinct promoters, LAMA3 generates two structurally distinct LMs; a so-called “full-length” variant LMα3b, and the much shorter LMα3a [8, 13–15]. The LaNt proteins are also derived from LM-encoding genes, through intron-retention and polyadenylation within the retained intron [11]. Four LaNt family members have been identified at the transcript level; however, only LaNt α31, derived from the LAMA3 gene, has been confirmed at the protein level [11]. LaNt α31 displays widespread tissue distribution [11] and is enriched in structured regions of ECM surrounding terminal duct lobular units (TDLUs) in normal breast tissue [16].

LaNt α31 functions have only been studied in corneal and skin epithelium to date, where upregulation of LaNt α31 was observed in response to corneal burn wounds or stem cells activation in ex vivo models, and where knockdown in expression reduced the rate at which epidermal keratinocytes close scratch wounds [11, 12]. Mechanistic studies have also indicated a role for this protein in modifying cell adhesion and migration via changes to matrix organisation and adhesion complex maturation [12, 17]. Further indications to LaNt α31 function come from its structure. Although LaNt α31 is smaller than LMs and lacks the coiled-coil domain required for LM trimer formation, it does share structural domains with LMα3b. Specifically, LaNt α31 is comprised a LM N-terminal domain (LN domain) and two LM-type epidermal growth factor-like repeats (LE domains) [11]. LN domains are involved in LM-to-LM interaction, and therefore are essential for laminin network assembly in BMs [7, 18, 19]. LaNt α31 also contains 54 unique amino acids with no homology to known structural motifs but which have allowed specific antibodies to be raised against this protein [11, 12]. Intriguingly the LaNt α31 protein architecture is structurally similar to other members of the laminin superfamily family, the netrins. Netrins are predominantly known as signalling proteins; however, netrin-4, via its LN domain, can disrupt LM-LM interactions and change the structural characteristics of LM networks [20–22].

While functional data suggest that LaNt α31 could be capable of influencing tumour progression, further rationale for investigating this protein in tumour microenvironment comes from studies of the other, more comprehensively studied, products of the LAMA3 gene. Reduction of LMα3a and LMα3b in breast carcinoma has been independently reported by several groups [23–25], with LMα3b downregulation in tumour vasculature associated with later stage tumours [26]. However, the situation is more complicated than a simple linear relationship, as increased LMα3 has been associated with triple negative breast carcinoma, and increased immunoreactivity that correlated with tumour stage has also been reported with antibodies against conformational epitopes in LMα3β3γ2 (LM332), LMβ3 and LMγ2, the preferred trimerization partners of LMα3a and LMα3b [25, 27].

Here we performed the first investigation into LaNt α31 in breast cancer. Breast tissue from normal and invasive ductal carcinomas were processed for immunohistochemistry with antibodies against LaNt α31, and correlation between staining intensity and pathology determined. LaNt α31 expression was upregulated in cultured breast carcinoma cells in culture and the impact on cellular behaviour was determined. The results revealed that LaNt α31 is increased in tumour tissue and that increased expression changes the mode of invasion of carcinoma cells into LM-rich matrices.

## Methods

### Ethical approval

The Liverpool Bio-Innovation Hub Biobank conferred ethical approval in writing for the use of samples in this project (REC reference 14/NW/1212, NRES Committee North West – Haydock). Project specific ethical approval for working with human tissue was conferred in writing by the University of Liverpool Research Ethics Committees (approval number:7488).

### Antibodies

Mouse monoclonal antibodies raised against human LaNt α31 were described previously [12, 16] and were used at 0.225 μg mL^−1^ for IHC and 1.8 μg mL^−1^ for immunoblotting. Mouse monoclonal antibodies against human LMα3 (clone CL3112) and mouse IgG (both Sigma-Aldrich, St. Louis, Missouri, USA) were used at 0.5 μg mL^−1^.

### Immunohistochemistry

Pilot tissues were obtained from the Liverpool Bio-Innovation Hub Biobank, all other TMA sections were purchased from Reveal Bioscience (product codes: BC02, BC03, BC05, BC06, and BC10; Reveal Bioscience, San Diego, USA) or US Biomax (product code: HBreD145Su02; US Biomax, Rockville, Maryland, USA). Sections were dewaxed and processed using a Leica Bond autostainer with Bond™ Polymer Refine Detection system (Leica Biosystems, Wetzlar, Germany). Briefly, following dewaxing, antigen retrieval was performed by incubating with a Tris/EDTA (pH 9 solution) solution for 20 mins at 60 °C, then endogenous peroxidases were blocked for 5 mins at room temperature with Bond hydrogen peroxide solution. Sections were incubated with primary or isotype-matched control antibodies at room temperature for 30 mins in Bond primary Ab solution (Tris-buffered saline, TBS, containing surfactant and protein stabilizer), then secondary anti-mouse IgG antibodies (<10 μg mL^−1^) with 10 % v/v animal serum in TBS were added for 15 mins at room temperature. DAB (66 mM) chromogen substrate was added for 20 mins at room temperature, and counterstaining performed with 0.1 % w/v haematoxylin for 5 mins. At each stage, washing was performed with Bond wash solution (TBS containing surfactant). Sections were finally dehydrated through a series of ascending ethanol concentration and then mounted with Pertex (all reagents Leica Biosystems). Stained tissue sections were imaged on the Aperio ImageScope slide scanner and processed using ImageScope software (Leica Biosystems).

### Immunohistochemistry interpretation

TMA cores were graded from 0-3 based on LaNt α31 immunoreactivity. Scores of 0 and 1 were then combined, and expression defined as low, medium, or high. All cores were scored by three independent scorers, and the mean score from duplicate cores used in final analyses. All patient data, including tumour/ node/ metastasis (TNM) status, tumour grade, and IHC marker scores (antigen ki-67 [ki67], epidermal growth factor receptor [EGFR], human epidermal growth factor receptor 2 [Her2], oestrogen receptor [ER], and progesterone receptor [PR]) were provided by Reveal Biosciences. Data were rounded to the nearest integer for intensity scores where required. For Ki67 percentage cell staining, scores were grouped as 0, 6, 6-10, or >10%, as provided by Reveal Biosciences.

### Cell culture

MCF-7 [28] and MDA-MB-231 [29] cells were cultured in high glucose (4.5 g L^−1^) Dulbecco’s Modified Eagle Medium (DMEM) (Sigma-Aldrich) supplemented with 10 % Foetal Calf Serum (FCS, LabTech International Ltd, Heathfield, East Sussex, UK) and 4 mM L-glutamine (Sigma-Aldrich).

### LaNt α31 expression

Full length *LAMA3LN1-eGFP* and *eGFP* adenoviral particles were prepared and used as previously described [12]. Transduction efficiency was determined by live fluorescent imaging at the time of analysis and expression confirmed by immunoblotting after 24 hours. Cells were homogenized by scraping into urea/ sodium dodecyl sulphate (SDS) buffer (10 mM Tris-HCl pH 6.8, 6.7 M Urea, 1 % w/v SDS, 10 % v/v Glycerol and 7.4 μM bromophenol blue, containing 50 μM phenylmethysulfonyl fluoride and 50 μM N-methylmaleimide, all Sigma-Aldrich). Lysates were sonicated and 10 % v/v β-mercaptoethanol added (Sigma-Aldrich). Proteins were separated by SDS-polyacrylamide gel electrophoresis (SDS-PAGE) using 10 % polyacrylamide gels (1.5 M Tris, 0.4 % w/v SDS, 10 % acrylamide/ bis-acrylamide; electrophoresis buffer; 25 mM Tris, 190 mM glycine, 0.1 % w/v SDS, pH 8.5 all Sigma-Aldrich). Proteins were transferred to nitrocellulose membranes (Biorad, California, USA) using the Biorad TurboBlot™ system and blocked for one hour at room temperature in Odyssey^®^ TBS-Blocking Buffer (Li-Cor BioSciences, Lincoln, Nebraska, USA). The blocked membranes were incubated overnight at 4°C with primary antibodies diluted in blocking buffer, then probed for 1 hour at room temperature with IRDye^®^ conjugated secondary antibodies against mouse IgG (800CW) raised in goat (LiCor BioSciences) diluted in Odyssey^®^ TBS-Blocking Buffer buffer at 0.05 μg mL^−1^. Membranes were imaged using the Odyssey^®^ CLX 9120 infrared imaging system and Image Studio Light v.5.2 (LiCor BioSciences) used to process scanned membranes.

### Resazurin reduction assay

Transduced or non-transduced cells were plated in triplicate at 1.5 x 10^4^ cells/well of a 96-well plate (Greiner-Bio One, Kremsmunster, Austria). After 24 hours culture media was replaced with serum-free and phenol red-free media containing 44 μM resazurin sodium salt (Sigma-Aldrich) [30]. The cells were then incubated for 2 hours. The media was removed and transferred to a black 96-well plate (Greiner-Bio One) and fluorescence measured at 570 nm using a SPECTROstar plate reader (BMG LABTECH, Ortenberg, Germany).

### 2D migration assays

For gap closure assays, cells were seeded into ibidi^®^ 2-well culture inserts (ibidi, Martinsried, Germany); at 7.0 x 10^4^ cells/well (MCF-7) or 8.0 x 10^4^ cells/well (MDA-MB-231). Culture inserts were carefully removed after 6 hours, cell debris washed away, and the gap margin imaged using brightfield optics on a Nikon TiE epifluorescence microscope with a 10X objective at 0 and 16 hours (Nikon, Tokyo, Japan). Gap closure was measured as a percentage relative to starting area using the freehand tool in image J (NIH, Bethesda, MA).

For low-density migration assays, cells were seeded at 5.0 x 10^4^ cells/well of a 12-well plate, then imaged every 2 minutes over a 2-hour period using a 20X objective on a Nikon Eclipse Ti-E fluorescent microscope adapted for live cell imaging. Individual cells were tracked using the MTrackJ plugin on image J and migration speed calculated.

### Inverted invasion assay

Inverted invasion assays were performed as previously described [31–33]. Briefly, 100 μL of 1 mg mL^−1^ rat tail collagen I (Corning Inc., New York, USA) or 4 mg mL^−1^ Matrigel^®^ from Engelbreth-Holm-Swarm (EHS) mouse sarcoma [34], was pipetted on top of Transwell^®^ 24-well, 0.8 μm, polycarbonate inserts (Corning). Then collagen was gelled through addition of 9.2 mM NaOH. Once gelled, the inserts were inverted and 100 μL of cell suspension containing 8.0 x 10^5^ cells were added to the lower surface. The transwells were than incubated for 4 hours to allow the cells to attach before returning to the original position with basal side downward. 1 mL of serum-free media was pipetted into the lower chamber of the transwell, and 100 μL of normal culture media supplemented with 25 ng mL^−1^ EGF, as a chemoattractant, was added to the upper chamber [35]. After 72 hours, the cells were fixed with 3.7 % v/v formaldehyde for 30 mins, permeabilised with 0.05 % v/v Triton X-100 then stained for 1 hour with DAPI (Sigma-Aldrich). The inserts were mounted onto a glass coverslip and imaged with a Zeiss Marianas (3i) spinning-disk confocal microscope by taking a z-stack with images every 5 μm, using SlideBook 5.5 software (3i, Intelligent Imaging Innovations Ltd, London, UK).

An algorithm was generated to automatically measure the DAPI stained nuclei in each slice of the z-stack. The average nuclei size was established by taking a range of manual measurements through the different planes of the z-stack. The average was used to set: i) the intensity threshold for distinguishing nuclei fluorescence from background as *t=1000*, ii) the expected nuclei size *s=240* pixels, corresponding to an area of 101.4 μm or a radius of 5.68 μm, ii) the lower size threshold *s^-=120* pixels (area = 50.7 μm or a= radius of 4.02 μm), below which captured objects were not considered to be nuclei and iv) the upper size threshold *s^Λ^+ = 400* pixels (area 169 μm or radius of 7.33 μm), above which the captured object was assumed to be an artefact. For each image slice, the image was imported and a coarse segmentation of the nuclei performed by thresholding at the intensity value *t* to achieve a binary nuclei/non-nuclei image. Any nuclei within an absolute distance of 0.65 μm or 1 pixel were connected to take into account noise, and an initial index taken of the distinct identified nuclei candidates, while measuring the size of each nuclei.

Individual cells were then identified as follows: i) any identified object under *s*^-, *s/2* pixels or over *s*^+ pixels in size was considered to be noise, ii) any captured object within the thresholds *s/2* and *3s/2* was considered to be one nuclei, iii) any remaining captured object which was larger than *3s/2* in size was considered to be a cluster of nuclei which could not be split due to the resolution of the image. In these cases, the number of cells in the cluster was calculated as (*cluster size) /s*, rounded to the nearest integer. The following measures were then taken: i) total luminance of the cell image, ii) PixCount: the number of pixels considered to be a cell after thresholding, corresponding to the total area in microns of the region considered to contain a cell. iii) Cell Count: the total number of cells identified, including those estimated from cell clusters, iv) Entropy: a measure of randomness of the thresholding data, which identified how clustered the cells in the image were.

### Data analyses

Microsoft Excel (Microsoft, Washington. USA), Graphpad Prism v.6 (Graphpad Software, California, USA), or SPSS statistic 24 (IBM Corporation, New York, USA) were used to analyse numerical data and generate graphs. Figures were generated using CoralDraw 2017.The Wilcoxon signed-rank test or Somers’ D was used for ordinal data, or Mantel-Cox log-rank test for survival data in immunohistochemistry analyses, and one-way ANOVA with Bonferroni post hoc test were used for continuous variables. Differences were deemed statistically significant where type I error rates were below 5 %.

## Results

### LaNt α31 and LMα3 display distinct distribution patterns in invasive ductal carcinomas

First we compared the distribution of LaNt α31 and LMα3 in a pilot panel of four normal (Figure 1A), four invasive ductal (Figure 1B) and four triple negative (Er-PR-Her2-) invasive ductal tissue sections (Figure 1C). The LMα3 antibodies used recognise both the LMα3a and LMα3b forms. LaNt α31 and LMα3 displayed very similar distribution in the normal tissue (Figure 1A), primarily restricted to TDLU as previously reported [16]. However, much stronger and more widespread LaNt α31 expression was detected in three of the four invasive ductal and three of four ER-, PR-, Her2-specimens. In the invasive ductal tissue, LaNt α31 displayed a more widespread distribution than LMα3 (arrowheads, Figure 1B), whereas in ER-PR-Her2-cases the LMα3 and LaNt α31 distribution were very similar (Figure 1C). LMα3 has been extensively investigated in breast cancer [23–25]; however, these new data for LaNt α31 indicated potential additional value of investigating this isoform independently from LMα3 in invasive ductal carcinoma.

**Figure 1:**
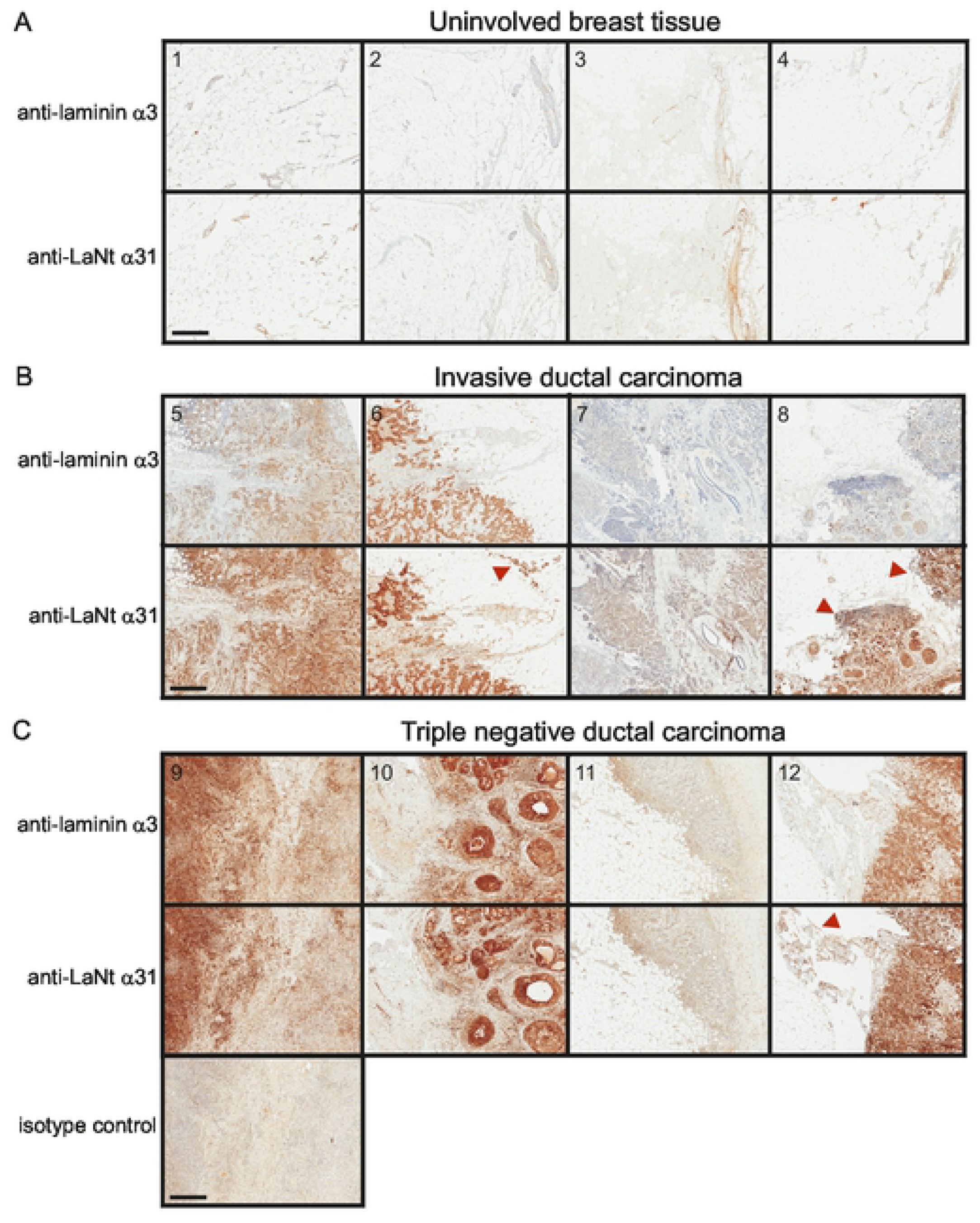
LaNt α31 is upregulated in ductal carcinoma. Serial sections from formalin-fixed paraffin-embedded human breast tissue processed for immunohistochemistry with mouse monoclonal antibodies against laminin α3, LaNt α31, or mouse IgG-isotype control, (A) uninvolved breast tissue, (B) invasive ductal carcinoma, and (C) ER-, PR-, Her-invasive ductal carcinoma. Scale bars: 500 μm.

### LaNt α31 expression is elevated in invasive ductal carcinoma and in nodal metastases compared to primary tumour tissue

To formally determine whether LaNt α31 expression levels change in invasive ductal carcinoma, the relative intensity of cellular immunoreactivity in epithelial-like tissue was compared between normal breast tissue compared with tumour biopsies from the same person, with intensity scored by three independent, blinded scorers (representative examples, Figure 2A, all cores Figure S1A, patient demographics Table 1). These paired analyses revealed LaNt α31 expression to be increased in the cancer specimen in 14 of 25 tumours (56%), eight had no change, while three displayed decreased expression in the cancer tissue (Wilcoxon signed ranks test, z=-2.67, p=0.008, Figure 2A).

**Figure 2:**
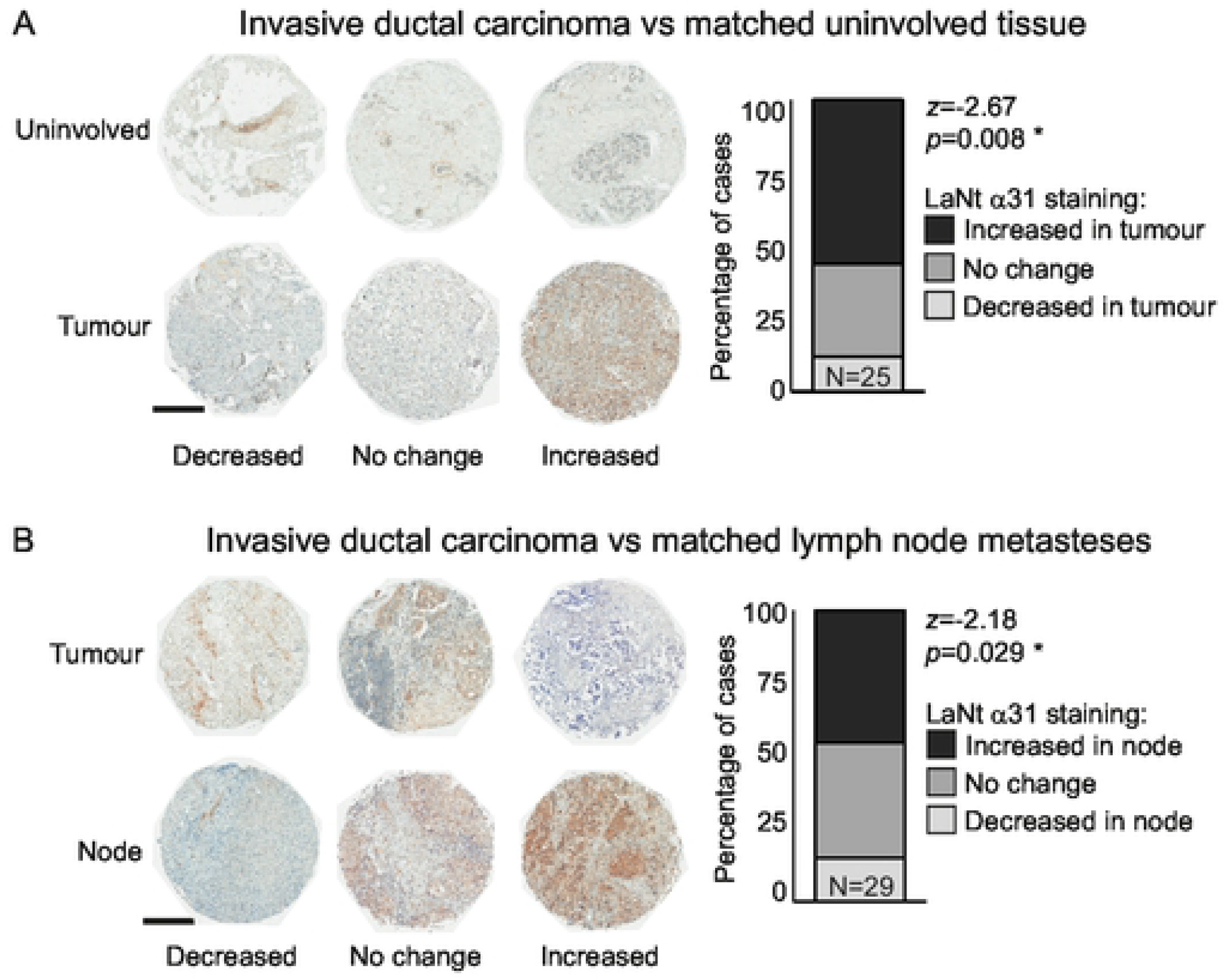
LaNt α31 is upregulated in ductal carcinoma and in lymph node metastases. Formalin-fixed paraffin-embedded human breast tissue microarray sections processed for immunohistochemistry with mouse monoclonal antibodies against LaNt α31. Two separate arrays were used; (A) uninvolved with paired invasive/ in situ ductal carcinoma tissues (N=25), (B) invasive ductal carcinoma with paired node metastases (N=29). Cores were scored as either decreased, no change, or increased staining intensity relative to the paired uninvolved (A) or primary tumour (B) core from the same donor (representative images shown). Stacked columns of percentage of cases in each category were plotted and Wilcoxon signed ranks test used to describe observed relationship. Scale bars: 500 μm.

**Table 1-.** Patient ages for each study cohort

| Study cohort | Mean | Median | Range | N |
| --- | --- | --- | --- | --- |
| Invasive ductal carcinoma vs uninvolved | 46 | 45 | 29-70 | 25 |
| Invasive ductal carcinoma vs lymph node metastases | 47 | 46 | 30-69 | 29 |
| Ductal carcinoma | 46 | 44 | 21-86 | 166 |
| Invasive ductal carcinoma | 60 | 57 | 33-88 | 132 |

Comparison of staining intensity between cores taken from the primary tumour against those from nodal metastasis from the same person using the same scoring approach, revealed that 13 of the 29 (45%) of the nodal metastasis displayed stronger LaNt α31 staining compared with primary tumour tissue, 12 were scored the same, and four were decreased in the nodal tissue (Figure 2B, Figure S1B, patient demographics Table 1, z=-2.18, p=0.029).

### LaNt α31 upregulation is associated with less proliferative tumours

To determine if LaNt α31 immunoreactivity held prognostic value for invasive ductal carcinoma, we processed 166 patients (Table 1) with two cores scored per patient by three independent scorers with the scores defined as “low”, “medium” or “high” LaNt α31 intensity (Figure 3A). First, we asked if LaNt α31 staining intensity was predictive for tumour grade, Ki67 expression (i.e. high proliferation [36, 37]), nodal involvement, or survival (Figure 3B-E, respectively). For tumour grade, although an increased proportion of cores displayed strong LaNt α31 staining in higher grade tumours, this association did not reach statistical significance (d=0.109, p=0.146, Figure 3B). Surprisingly, a negative correlation was observed between Ki67 expression level and LaNt α31 staining intensity (d=-0.3, p=0.006, Figure 3C). The relative proportions of high, medium and low LaNt α31 expression were broadly similar between those cases which presented with nodal involvement and those without (Somers’ d=0.059, p=0.505, Figure 3D).

**Figure 3:**
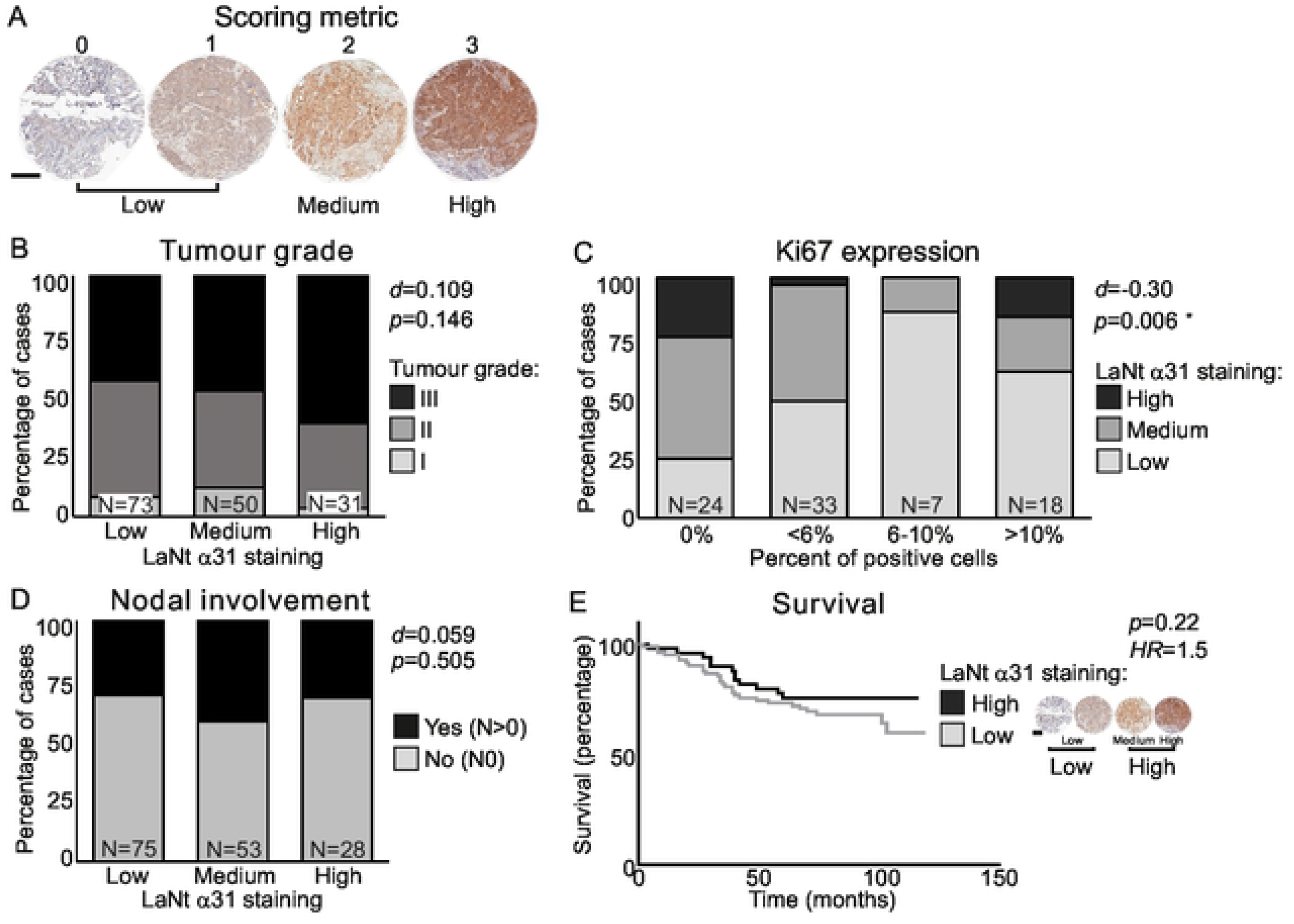
LaNt α31 upregulation in ductal carcinoma does not correlate with nodal involvement or tumour grade. Formalin-fixed paraffin embedded human breast tissue microarray sections processed for immunohistochemistry with mouse monoclonal antibodies against LaNt α31. Cores were scored based on LaNt α31 staining intensity from 0-3. Scores of 0 or 1 were combined and designated as low LaNt α31 expression, score 2 as medium expression, and 3 as high expression. (A) Representative example of core scoring. (B-D) Stacked column graphs of percentage of cases with each staining intensity segregated by tumour grade I, II, or III (B), Ki67 expression (C), or by nodal involvement (D). Somers’ D was used to describe observed relationships between LaNt α31 staining intensity and the independent variables. (E) Kaplan–Meier survival curve, were LaNt α31 staining intensity was simplified to low or high by pooling medium and high cores. Logrank was used to determine hazard ratio and chi square for significance. Scale bar in (A): 300 μm.

Survival was assessed for the 132 cases where these data were available. To account for the smaller sample size here, LaNt α31 staining intensity was simplified to either low or high expression by combining the medium with the high expression cases. These data revealed that tumours with higher LaNt α31 staining intensity had marginally improved overall survival, although this did not reach statistical significance at an α of 0.05 (low=66% survival, high=76% survival, hazard ratio 1.52, confidence interval 0.8-2.8, p=0.22, Figure 3E).

Next, we asked if LaNt α31 staining displayed a relationship with any of the commonly used breast cancer biomarkers (Figure 4). These data revealed a positive association between LaNt α31 and EGFR (EGFR d=0.26, p=0.003, Figure 4A), while observed associations with Her2, ER and PR were each below statistical significance at an α of 0.05 (Figure 4B-D). However, the data suggest a potential weak, negative association with ER (d=-0.12, p=0.056).

**Figure 4:**
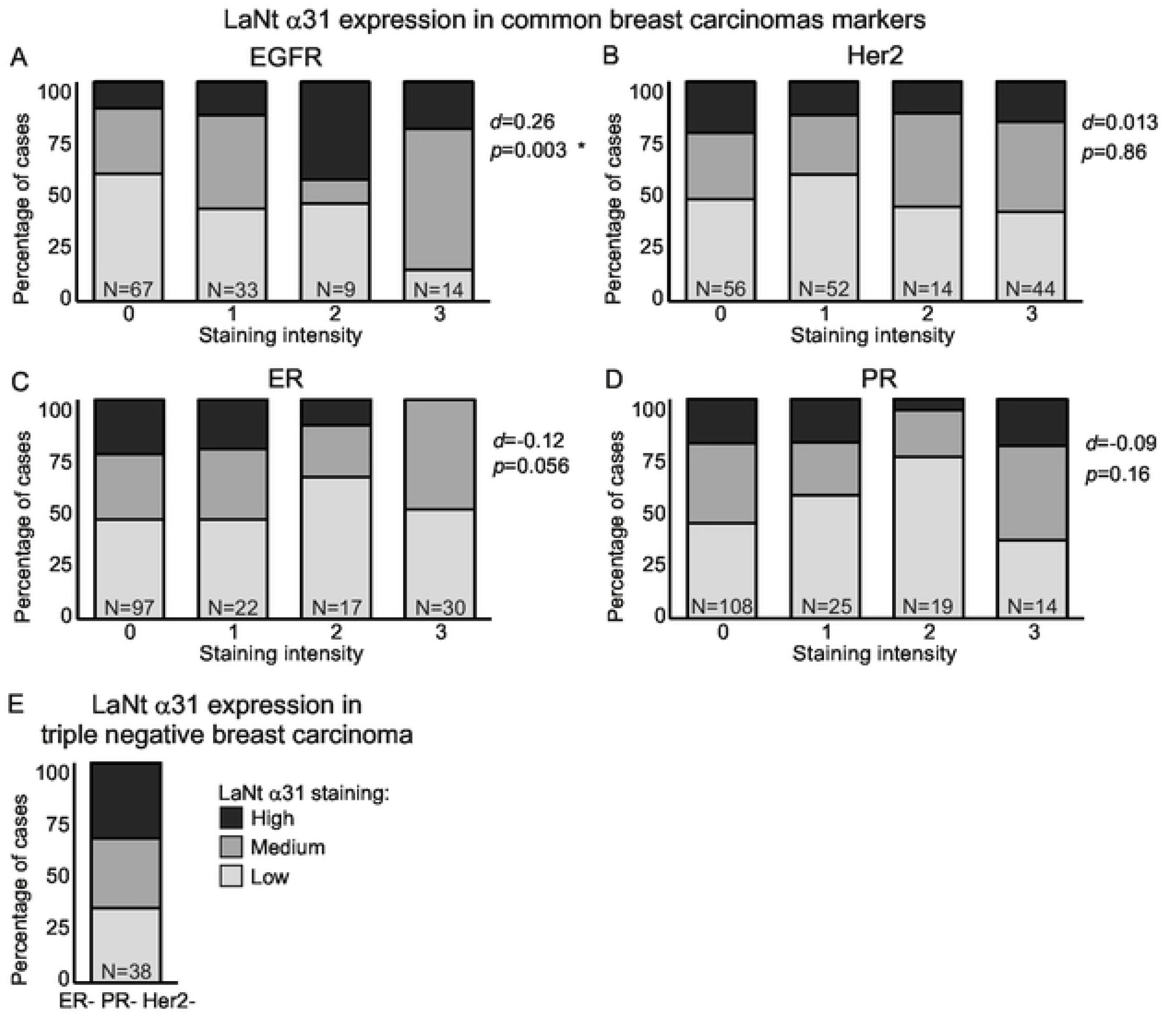
Positive correlation between LaNt α31 staining intensity and EGFR expression. Formalin-fixed paraffin embedded human breast tissue microarray sections processed for immunohistochemistry with mouse monoclonal antibodies against LaNt α31. Cores were scored based on LaNt α31 staining intensity from 0-3. Scores of 0 or 1 were combined and designated as low LaNt α31 expression, scores 2 as medium expression, and 3 as high expression. Stacked column graph of percentage of cases that fall into group after segregation based on pathologist provided grading of immunohistochemistry markers; (A) EGFR, (B) Her2, (C) ER, (D) PR, or (E) ER- PR- Her2- cases. Somers’ D was used to describe observed relationship between LaNt α31 staining intensity and independent variable.

The cores from triple negative cancers were analysed separately (Figure 4E, n=38 cores). These data revealed that more than 65 % of the ER- PR- Her2- cores had either medium or high LaNt α31 staining intensity.

### LaNt α31 expression does not induce increased 2D migration or invasion in a non-motile cell line

Together the immunohistochemistry findings indicate an association between LaNt α31 staining and a subset of tumours; however, this does not necessarily indicate that expression change contributes to the disease. To address this question, we used an adenoviral system to drive overexpression of LaNt α31 tagged with eGFP (+LaNt α31- eGFP) in two widely used cell lines derived from invasive ductal carcinomas; MCF-7 and MDA-MB-231 (Figure S2A-B). Cells transduced with eGFP only were used to control for adenoviral transduction and eGFP expression. Expression of eGFP or LaNt α31-eGFP had no effect on metabolic activity of either MCF-7 or MDA-MB-231 cells (Figure S2C-D).

MCF-7 cells migrate slowly and do not invade in 3D invasion assays [38–40]. This line was therefore used to assess if induction of LaNt α31 expression was sufficient to increase these behaviours. In gap closure assays (Figure S3A-B), MCF-7 cells expressing LaNt α31 closed the gap more slowly than controls in three out of seven independent experiments; however, overall the differences were not statistically significant (Figure S3B median: MCF-7 83.3 %, +eGFP 79.7 %, +LaNt α31-eGFP 67.8 %, *p*=0.37). In single cell assays (Figure S3C-D), a small increase in migration speed was observed three out of four independent experiments, although differences between LaNt α31 and control treatments were not statistically significant (mean migration speed +/- S.D. MCF-7 0.31 μm min^−1^ +/- 0.13, +eGFP 0.36 μm min^−1^ +/- 0.09, +LaNt α31-eGFP 0.35 μm min^−1^ +/- 0.06, *p*=0.53, determined by ANOVA. Figure S3D).

Next, we asked if increasing LaNt α31 expression could induce invasive capabilities in MCF7 using an inverted invasion assay [33, 41], where cells were seeded on the base of a porous membrane then stimulated to invade into a provided matrix and using an EGF gradient as a chemoattractant (Figure S3E-H). As LaNt α31 effects could be LM specific, invasion into two different matrices were analysed; collagen I to mimic the interstitial matrix (Figure S3E-F) and Matrigel, a BM analogue that contains approximately 60 % LM111, 30 % Type IV collagen, and 8 % entactin [34, 42] (Figure S3G-H). As expected, untreated MCF-7 cells had very low invasive capabilities into both matrices, and overexpression of LaNt α31 did not lead to any significant change in invasion depth (mean +/- S.D invasion depth into collagen I: MCF-7 57 μm +/- 24, +eGFP 55 μm +/- 22, +LaNt α31-eGFP 50 μm +/- 17, *p*=0.78, determined by ANOVA. Figure S3F. Into Matrigel: MCF-7 55 μm +/- 0, +eGFP 57 μm +/- 13, +LaNt α31-eGFP 58 μm +/- 6, *p*>0.71, determined by ANOVA. Figure S3H).

### Increased LaNt α31 expression causes a change in mode of invasion into laminin-rich hydrogels

In contrast to MCF7, MDA-MB-231 are much more motile and invasive [33, 41, 43]. However, the effect of LaNt α31 expression on the 2D migration of MDA-MB-231 was minimal, with a slight, but not statistically significant reduction in gap closure rate (median: MDA-MB-231 98.8 %, +eGFP 92.7 %, +LaNt α31-eGFP 85.1 %, *p*=0.23. Figure 5A-B), and similar outcome in single cell migration assays (mean speed +/- S.D: MDA-MB-231 0.48 μm min^−1^ +/- 0.10, +eGFP 0.50 μm min^−1^ +/- 0.14, +LaNt α31-eGFP 0.36 μm min^−1^ +/- 0.15, MDA-MB-231 vs +LaNt α31-eGFP, *p*=0.16; +eFGP vs +LaNt α31-eGFP, *p*=0.02 determined by ANOVA with Bonferoni post hoc test. Figure 5C-D).

**Figure 5:**
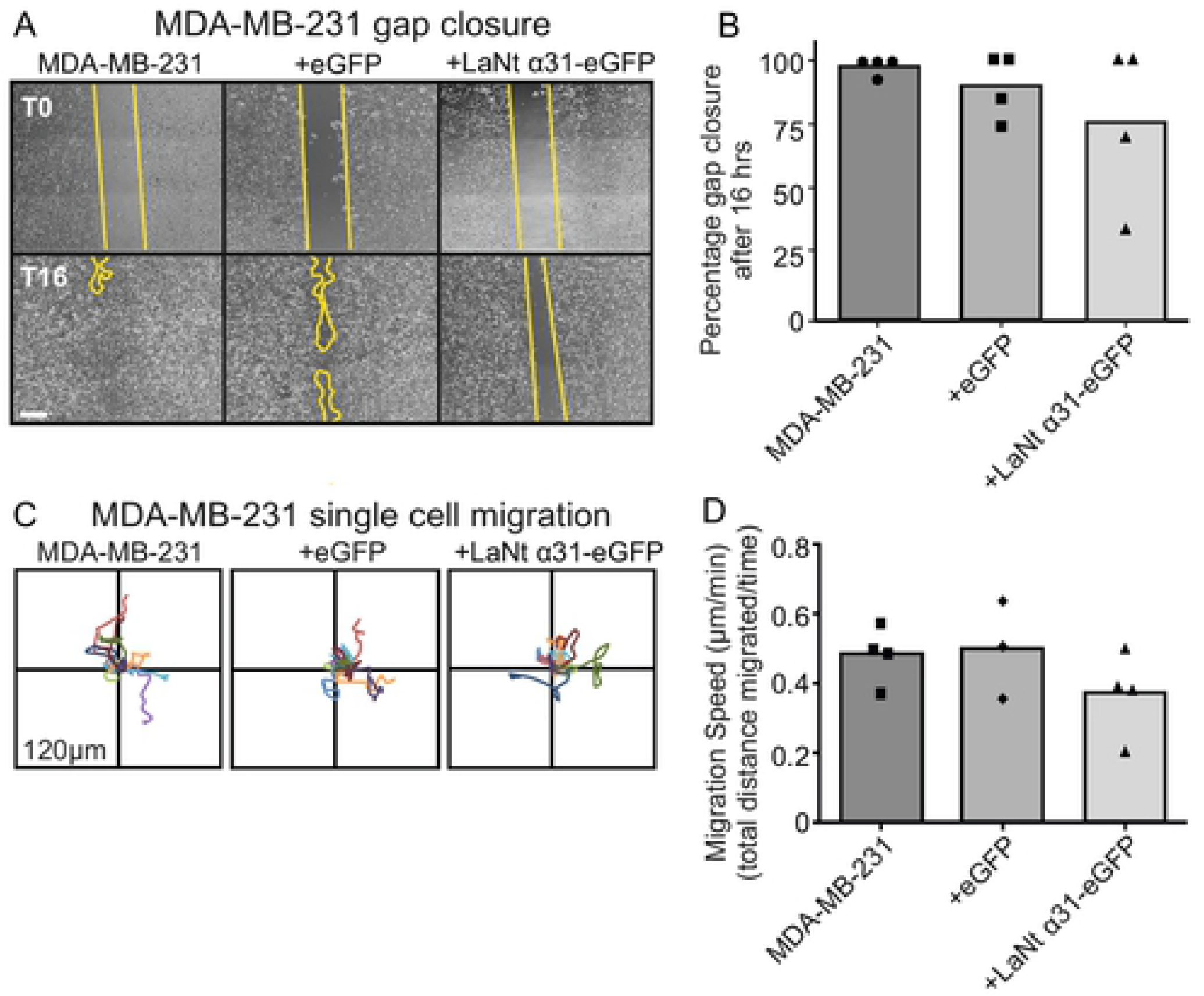
LaNt α31 overexpression does not significantly affect 2D migration MDA-MB-231 cells. MDA-MB-231 cells were either left untreated or transduced with eGFP (+eGFP), or LAMA3LN1-eGFP (+LaNt α31-eGFP). For gap closure assays, 24 hours after transduction, cells were seeded into ibidi^®^ 2-well culture inserts and allowed to attach for 6 hours, the inserts were then removed, and the gap margin imaged at 0 hours and 16 hours. For single cell migration assays, 24 hours after transduction, cells were seeded onto tissue culture plastic and the migration paths of individual cells tracked over a four-hour period. (A) Representative images from immediately after removing chamber (T0 upper panels) and after 16 hours (T16 lower panels), yellow lines delineate wound margins. (B) Gap closure was measured as a percentage relative to starting gap area. (C) Vector diagrams showing representative migration paths of 10 individual cells with each colour representing a single cell. (D) Migration speed was measured as total distance migrated over time. Each point on the associated dot plots represents an independent experiment with 2-3 technical replicates per experiment for gap closures assays or 20-40 cells per low density migration assay. Statistical tests of differences relative to controls were performed using one-way ANOVA followed by Bonferroni’s post hoc analyses; p>0.05 in all comparisons. Scale bar in (a) represents 100 μm.

As expected MDA-MB-231 invaded into both collagen I and Matrigel matrices (Figure 6). The invasion depth of +LaNt α31-eGFP MDA-MB-231 cells into collagen I was unchanged compared with controls; however, there was a slight reduction in total invasion depth into Matrigel although this did not reach statistical significance compared with both controls once adjusted for multiple comparisons (mean +/- S.D depth into collagen: MDA-MB-231 89 μm +/- 22, +eGFP 90 μm +/- 7, +LaNt α31-eGFP 87 μm +/- 18, *p*=0.52. Figure 6A-B. Invasion into Matrigel: MDA-MB-231 136 μm +/- 10 S.D, +eGFP 127 μm +/- 4, +LaNt α31-eGFP 97 μm +/- 11, MDA-MB-231 vs +LaNt α31-eGFP *p*=0.11, +eGFP vs +LaNt α31-eGFP *p*=0.14, ANOVA with Bonferoni post hoc test. Figure 6C-D).

**Figure 6:**
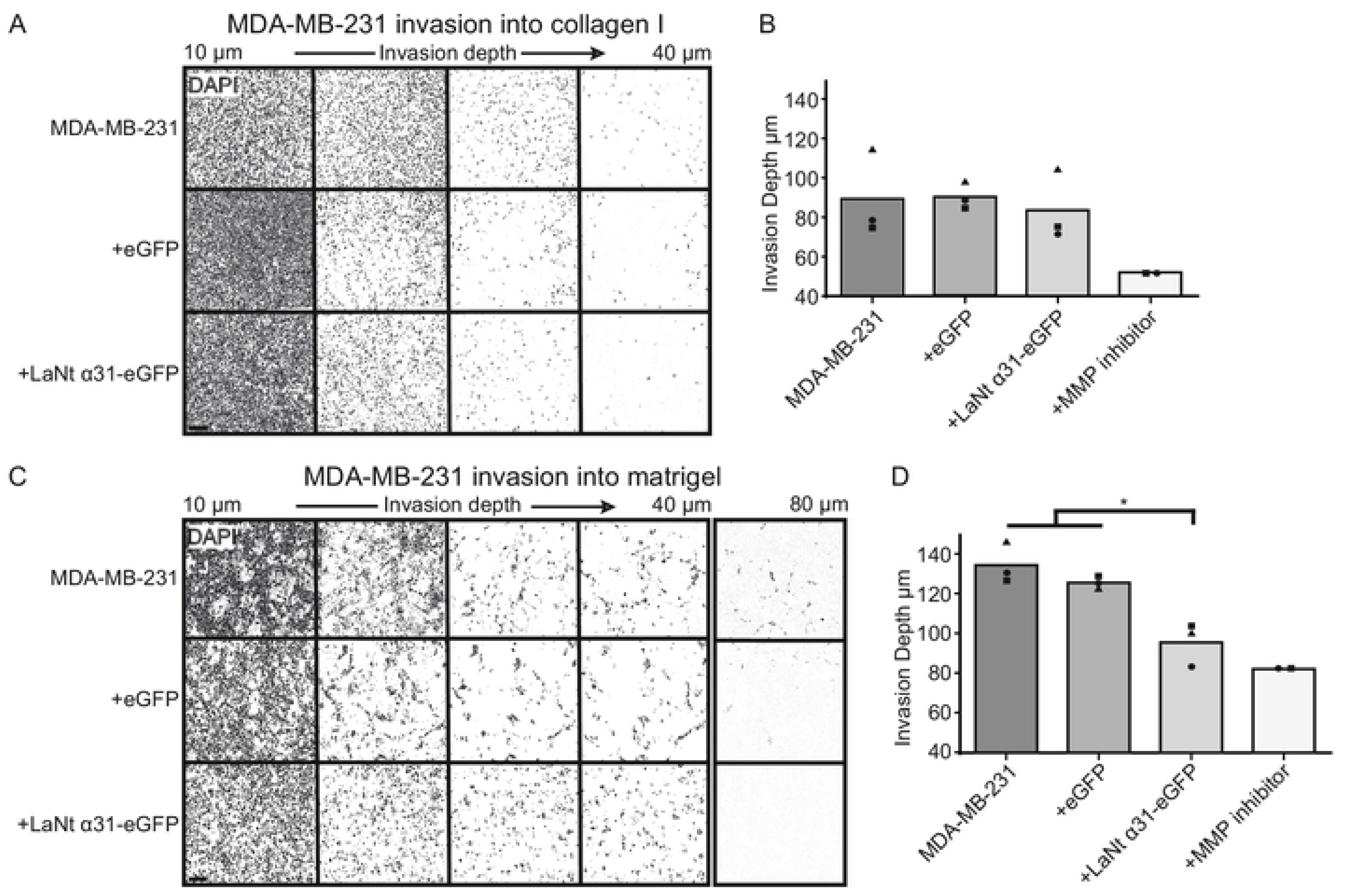
LaNt α31 overexpression causes a small reduction in invasion of MDA-MB-231 cells into Matrigel. MDA-MB-231 cells, either left untreated or transduced with eGFP (+eGFP), or LAMA3LN1-eGFP (+LaNt α31-eGFP), were plated onto the outside of a transwell membrane. 10 ng mL^−1^ epidermal growth factor was used to stimulate invasion through the membrane into collagen I (A-B) or Matrigel (C-D). After 48 hours, the cells were fixed and DAPI stained then imaged at 5 μm intervals using a spinning disk confocal microscope. (A) and (C) Representative images from 10-40 μm depth presented at equal intervals, with an additional slice at 80 μm in (C). Absolute invasion depth was measured where cell count ≥1. Treatment with GM6001 MMP inhibitor was included as an invasion inhibiting control. Each point on the graphs in (B) and (D) represents an independent experiment, with 2-3 technical replicates per assay. * represents p<0.05 between bracketed groups as determined by one-way ANOVA followed by Bonferroni’s post hoc analyses.

Although there were minimal differences in invasion depth, visual analysis of the Matrigel invasion assays, revealed an intriguing distinct phenotypic difference between the LaNt α31 overexpressing cells compared with the controls (Figure 6C and 7A). Whereas MDA-MB-231 cells usually invade into Matrigel as multicellular streams, as has been reported previously [43, 44], the +LaNt α31-eGFP cells did not display this behaviour (Movies 1 and 2, maximum intensity projection Figure 7A). To assess this quantitatively, we wrote a macro to convert the DAPI stained images at 60 μm depth into an inverse entropy score as measure of cohesiveness (entropy^−1^) (Figure 7B). At this invasion depth, cell densities were not different between +LaNt α31-eGFP and controls. These analyses agreed with our visual impression that the +LaNt α31-eGFP were less cohesive in their invasion profile (mean +/- S.D entropy^−1^ x10^−7^: MDA-MB-231 1.9 +/- 0.8, +eGFP 2.0 +/- 0.02, +LaNt α31-eGFP 0.93 +/-0.55, *p*=0.29. Figure 7C).

**Figure 7:**
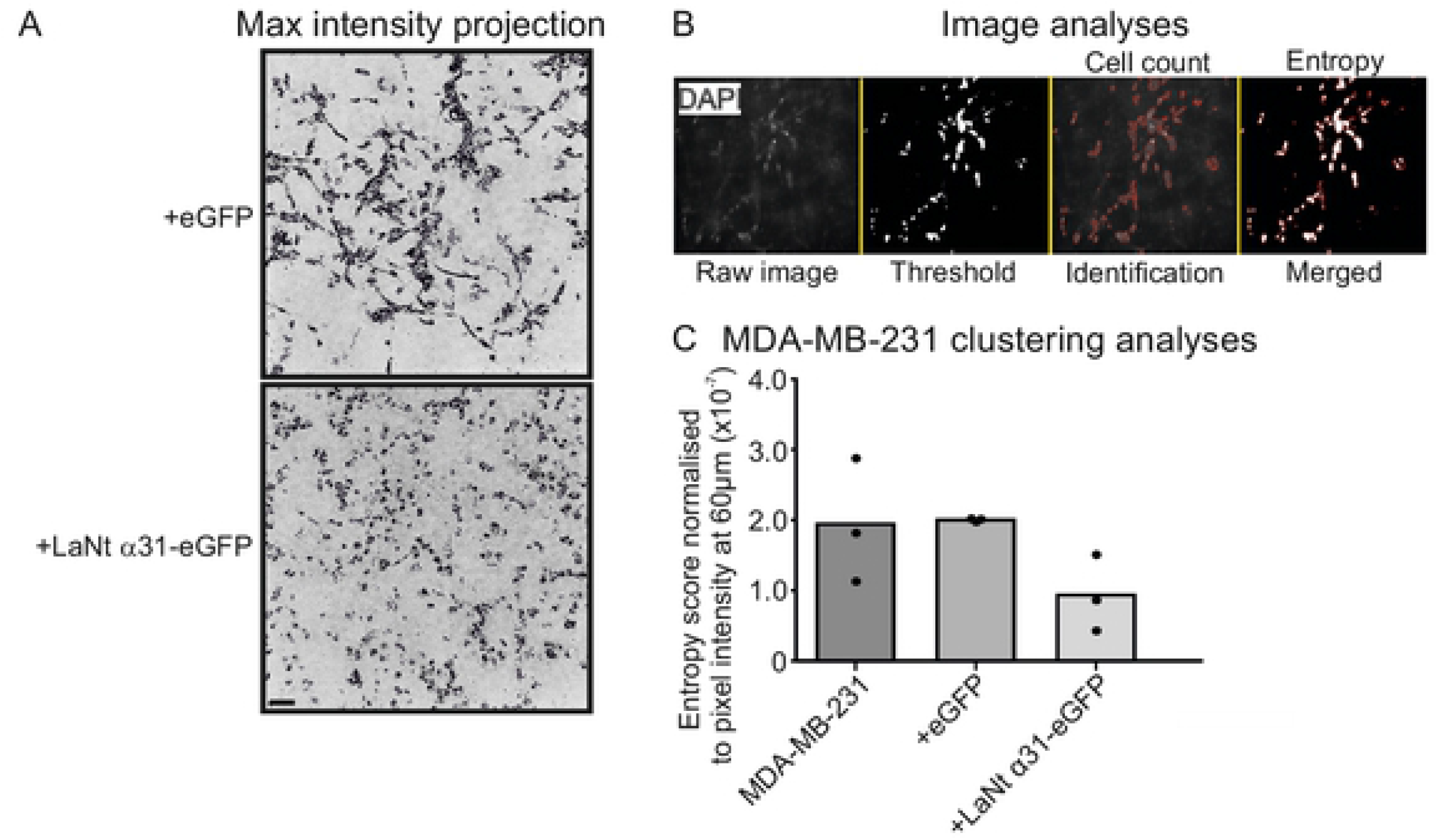
LaNt α31 overexpression causes a change in mode of invasion of MDA-MB-231 cells into Matrigel. MDA-MB-231 cells, either left untreated or transduced with eGFP (+eGFP), or LAMA3LN1-eGFP (+LaNt α31-eGFP), were plated onto the outside of a transwell membrane. 10 ng mL^−1^ epidermal growth factor was used to stimulate invasion through the membrane and into Matrigel. After 48 hours, the cells were fixed and stained with DAPI then imaged at 5 μm intervals using a spinning disk confocal microscope. (A) Maximum intensity projection of planes from 20-60 μm from the same assays in Figure 6C. (B) Image analyses method for determining entropy^−1^ as a measure of cell clustering; each stack of images was processed using an automated processing algorithm, where cell count and entropy score after a threshold was measured for each image in the stack. (C) Entropy^−1^ score at 60 μm normalised to pixel intensity plotted to assess clustering of cells.

### Tumours with high LaNt α31 expression are likely to be non-cohesive

As the LaNt α31 functional studies data suggested that high expression of this protein changes the mode of tumour invasion, we returned to the tissue array data, focusing specifically on the cores with high LaNt α31 intensity and assessed the tumour appearance in each of those cores as either “cohesive” or “non-cohesive” depending on whether tumour cells were present in contiguous islands with well-defined borders, (representative examples Figure 8A). These analyses revealed that 67.7 % of the high LaNt α31 expressing tumours were non-cohesive in appearance (21 of 31 cores, Figure 8B).

**Figure 8:**
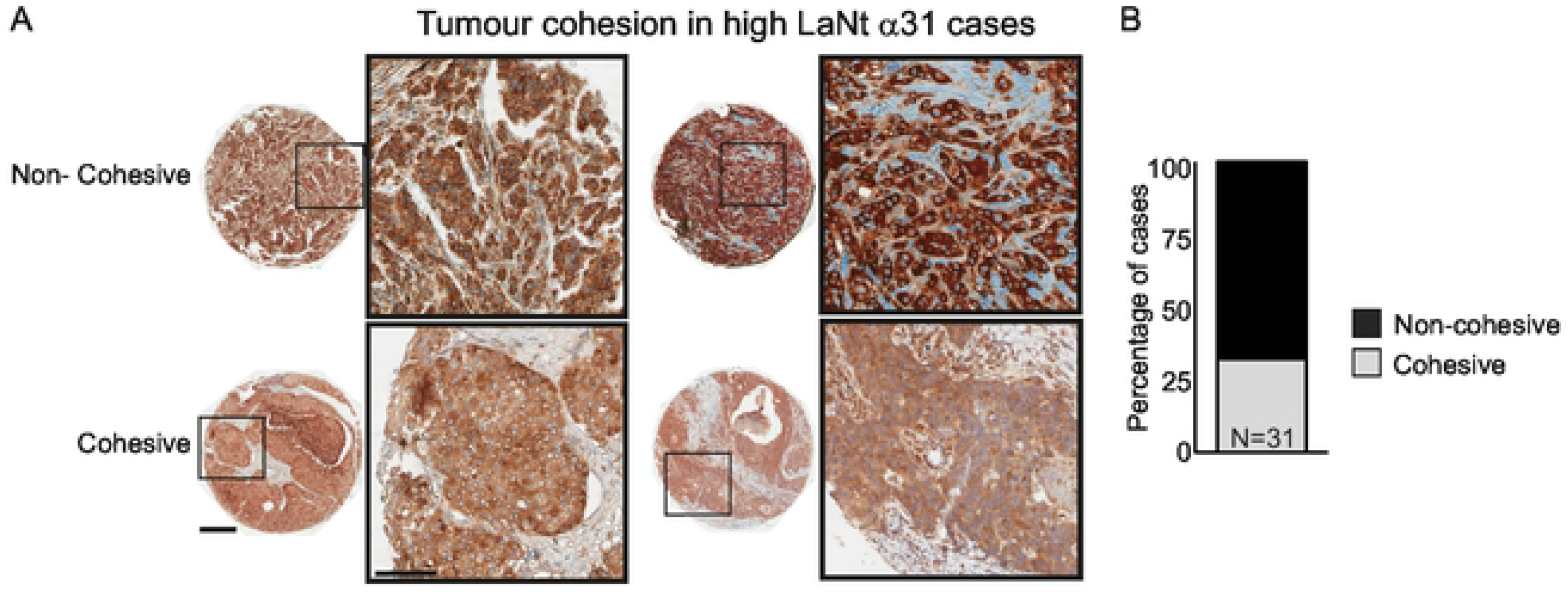
High LaNt α31 expression is associated with low tumour cohesion in invasive ductal carcinoma. Formalin-fixed paraffin embedded human breast tissue microarray sections processed for immunohistochemistry with mouse monoclonal antibodies against LaNt α31. Tumour cohesion was graded as either cohesive (tight tumour islands), or non-cohesive (chord-like) in tumour cores scored as having high LaNt α31 expression. (A) Representative example of core grading. (B) Stacked column graphs of percentage of cases that are either cohesive or non-cohesive. Scale bars: 300 μm.

## Discussion

The findings presented here have revealed that the little-known LM-related protein LaNt α31 is upregulated in a distinct sub-population of breast cancers, including a subset of TNBC, but is not definitively predictive of patient outcome. However, the results also show that LaNt α31 is capable of changing the mode of invasion of breast cancer cells into LM-rich matrices, suggesting that dysregulation of this protein could actively contribute to tumour pathology.

Usually in tumour situations, very few cells acquire the ability to invade and a hallmark of more aggressive tumours is plasticity in the modes of migration [45, 46]. A switch from multicellular streaming to individual cell invasion can happen in multiple overlapping ways. A major driver is the mechanical properties the ECM including matrix stiffness and the orientation of fibres; however decreased cell-cell adhesion, increased Rac-driven cytoskeletal protrusion compared with Rho-mediated contraction, an increased ability to generate traction forces, and differences in proteolytic activities all can drive the changes [43], as reviewed in [47]. Many of these mechanisms are intrinsically linked, which makes it challenging to directly assign a single behaviour to an individual protein. However, the Matrigel-specific effects and the structural similarity between netrin-4 and LaNt α31 make it tempting to predict that LaNt α31 has a disruptive effect on LM networks, softening and disordering the matrix through which the cells are invading [20, 22]. Matrigel contains many components beyond LM111 [42], therefore the observed phenotype could also be due to LaNt α31 interaction with these additional factors or acting through competition with cell surface receptors. LN domains, including the LN domain of LMα3b, are known to have cell-surface receptor-binding capabilities [48] [49], and notably netrin-4 and a proteolytically released LN domain fragment of LMβ1 are each capable of inducing epithelial to mesenchymal transition [20, 21, 50, 51]. Dissecting the mechanism of the LaNt α31-induced changes will not be a trivial undertaking but could be valuable to understand the process and as a route toward targeted intervention approaches.

An additional intriguing finding was the difference between LaNt α31 and LMα3. Although these proteins are genetically linked, they are structurally and functionally distinct. Specifically, LMα3a, as part of LM332, has been robustly demonstrated to enhance the migratory behaviour of MCF7 and MDA-MB-231 cells in culture [52, 53]. However, loss or focal disruption of LM332 staining is a more common feature in breast cancer, particularly for LMα3b, which shares a promoter with LaNt α31, and is downregulated in the tumour vasculature [26]. Indeed, in side-by-side comparison of the same tissue, the observation of different structures displaying upregulation of LaNt α31 compared with LMα3 points to differences in post-transcriptional regulation. We do not yet know if the difference is due to differences between the isoforms in terms of pre-mRNA processing, mRNA degradation, or post-translational proteolytic processing; however, these data do suggest that changes to LaNt α31 expression may have more widespread implications to other cancer subtypes where LMα3 is known to be dysregulated. In these contexts, processing tissue for LaNt α31 may have value as a biomarker.

## Conclusions

The combination of patient data and manipulative experimental data presented here have revealed LaNt α31, for the first time, to be associated with the progression of breast cancer. Moreover, the finding that LaNt α31 actively influences invasive behaviour indicate that targeting this protein’s function could hold potential as a therapeutic approach.

## Acknowledgements

This work was supported by Biotechnology and Biological Sciences Research Council Grants BB/L020513/1 and BB/P0257731, and by North West Cancer Research. These funding sources supported the purchase of consumables and equipment access and paid salaries of LDT.

LDT designed and conducted the experiments, analysed the data, and made the figures. TZ assisted in design, analysis and interpretation of the invasion assays. KJH designed the study, analysed data. All authors contributed to the writing and editing of the manuscript. The authors would also like to thank Bryan Williams for his help in writing image analysis macro, Abigail Pickett, Marian Jones, Eleanor Hughes, and Louisa Orfanou for their help in image scoring, and Louise Brown (funded by Breast Cancer Now grant 2014MayPR292) for help with the invasion assays. The authors would like to acknowledge the donors, for without whom, this work would not be possible.

## Abbreviations

(Ki67): Antigen ki-67
(BM): Basement membrane
(ER): Estrogen receptor
(ECM): Extracellular matrix
(EGF): Epidermal growth factor
(EGFR): Epidermal growth factor receptor
(eGFP): Enhanced green fluorescent protein
(Her2): Human epidermal growth factor receptor 2
(LaNt α31): Laminin N terminal protein α31
(LE-repeat): Laminin-type epidermal growth factor-like
(LM): Laminin
(LN domain): Laminin N-terminal
(p53): Tumour protein p53
(FFPE): Paraffin-fixed formalin-embedded
(PR): Progesterone receptor
(TDLU): Terminal duct lobular unit
(TNBC): Triple negative breast cancer

## Supporting information

**Figure S1: LaNt α31 is upregulated in ductal carcinoma and in lymph node metastases.** Formalin-fixed paraffin-embedded human breast tissue microarray sections processed for immunohistochemistry with mouse monoclonal antibodies against LaNt α31. Two separate arrays were used; (A) uninvolved with paired invasive/ in situ ductal carcinoma tissues (N=25), (B) invasive ductal carcinoma with paired node metastases (N=29). All paired cores analysed in Figure 2 ordered based on LaNt α31 staining intensity, low to high (left to right). Scale bars: 500 μm.

**Figure S2: LaNt α31 overexpression does not affect cell metabolism.** MCF-7 or MDA-MB-231 cells, either left untreated or transduced with eGFP (+eGFP), or LAMA3LN1-eGFP (+LaNt α31-eGFP), were maintained for 48 hours following transduction. (A) Immunoblots from total cell lysates for MCF-7 or MDA-MB-231 cells probed with antibodies against LaNt α31, with ponceau S total protein stained membrane below. (B) Resazurin salts (44 μM) were added to transduced cells for 2 hours, then the fluorescence intensity of the culture medium was measured at 570 nm. Dot plot of fluorescence intensity relative to non-transduced cells.

**Figure S3: LaNt α31 overexpression does not significantly affect 2D migration or 3D invasion of MCF-7 cells.** MCF-7 cells were either left untreated or transduced with eGFP (+eGFP), or LAMA3LN1-eGFP (+LaNt α31-eGFP). For gap closure assays, 24 hours after transduction, cells were seeded into ibidi^®^ 2-well culture inserts and allowed to attach for 6 hours, the inserts were then removed, and the gap margin imaged at 0 hours and 16 hours. For single cell migration assays, 24 hours after transduction, cells were seeded onto tissue culture plastic and the migration paths of individual cells tracked over a four-hour period. (A) Representative images from immediately after removing chamber (T0 upper panels) and after 16 hours (T16 lower panels), yellow lines delineate wound margins. (B) Gap closure was measured as a percentage relative to starting gap area. (C) Vector diagrams showing representative migration paths of 10 individual cells with each colour representing a single cell. (D) Migration speed was measured as total distance migrated over time. Each point on the associated dot plots represents an independent experiment with 2-3 technical replicates per experiment for gap closures assays or 20-40 cells per low density migration assay. For invasion assays, cells were plated onto the outside of a transwell membrane. 10 ng mL^−1^ epidermal growth factor was used to stimulate invasion through the membrane and into collagen I or Matrigel. After 48 hours, the cells were fixed and stained with DAPI then imaged at 5 μm intervals using a spinning disk confocal microscope. (E and G) Representative images of invasion into collagen I or Matrigel from 10-40 μm presented at equal intervals. (F and H) Absolute invasion depth was measured where cell count ≥1. Treatment with GM6001 MMP inhibitor was included as an invasion inhibiting control. Each point on the graphs represents an independent experiment, with 2-3 technical replicates per assay. * represents p<0.05 between bracketed groups as determined by one-way ANOVA followed by Bonferroni’s post hoc analyses. Statistical tests of differences relative to controls were performed using one-way ANOVA followed by Bonferroni’s post hoc analyses; p>0.05 in all comparisons. Scale bar in (a) represents 100 μm.

**Movie 1: MDA-MB-231 +eGFP invasion into Matrigel (.avi)**

MDA-MB-231 cells transduced with eGFP were plated onto the outside of a transwell membrane. 10 ng mL^−1^ epidermal growth factor was used to stimulate invasion through the membrane and into Matrigel. After 48 hours, the cells were fixed and stained with DAPI then imaged at 5 μm intervals using a spinning disk confocal microscope. Representative movie of invasion profile generated from z-stack slices for invasion between 20 and 60 μm after applying threshold.

**Movie 2: MDA-MB-231 +LaNt α31-eGFP invasion into Matrigel (.avi).** MDA-MB-231 cells transduced with LaNt α31-eGFP were plated onto the outside of a transwell membrane. 10 ng mL^−1^ epidermal growth factor was used to stimulate invasion through the membrane and into Matrigel. After 48 hours, the cells were fixed and stained with DAPI then imaged at 5 μm intervals using a spinning disk confocal microscope. Representative movie of invasion profile generated from z-stack slices for invasion between 20 and 60 μm after applying threshold.

